# Potassium stimulates fruit sugar accumulation by increasing carbon flow in Citrus

**DOI:** 10.1101/2023.11.06.565758

**Authors:** Kongjie Wu, Chengxiao Hu, Peiyu Liao, Yinlong Hu, Xuecheng Sun, Qiling Tan, Zhiyong Pan, Shoujun Xu, Zhihao Dong, Songwei Wu

## Abstract

Soluble sugar is a key factor of flavor quality in citrus. Potassium (K) is known as a quality element, which plays key roles in improving sugar accumulation and fruit quality, but the mechanisms are largely unknown. This study aims to elucidate how K improves sugar accumulation by regulating carbon flow between source and sink in Newhall navel orange (*Citrus sinensis*). The results demonstrated that appropriate K concentration improved fruit quality and sugar accumulations in citrus, and 1.5% of K concentration in pulp was the optimal concentration for fruit quality. K increased strength of sink and source, as supported by the increased fruit growth rate, gene expressions related to sugar metabolism and sugar transport in fruit, and photosynthesis, gene expressions of sugar metabolism in leaf, respectively, which contributed to increasing sugars accumulation. Further study indicated that K improved carbon flow from source to sink by boosting symplastic and apoplastic loading of phloem, as supported by the increased CF signal intensities, plasmodesmata densities, and the expressions of *CsSUT1* and *CsSUT2* in leaf at early and mid stages of fruit development, finally increasing sugar accumulation in fruit. Conclusively, K stimulates fruit sugar accumulation by increasing carbon flow in Newhall navel orange.

**Highlight:** K application stimulated carbon flow between source and sink through symplastic and apoplastic loading, which were supported by the structural characteristics of phloem and the expression of *CsSUTs* and *CsSWEET*s, ultimately promoting sugar accumulation in *Citrus sinensis*.

## Introduction

Potassium (K), as one of the macronutrients of plants, is essential for the growth and development of plants (Voitsekhovskaja et al., 2020). K is usually present in plants as K^+^ involved in numerous physiological processes, including regulation of photosynthesis, improvement of photoassimilate transport, increment of source carbohydrate synthesis, maintenance of cytoplasmic pH homeostasis (Reddy and Zhao, 2005; Oosterhuis et al., 2013; Wu et al., 2021). In citrus, which is a fruit of both special flavour and economical profitability, K application significantly increased soluble sugar accumulation by facilitating the assimilate transport and carbohydrate distribution and this increase was shown to be the main explanation of soluble sugars accumulation under appropriate K application conditions (Quaggio et al., 2006; Wu et al., 2021). However, the mechanism of K regulating assimilate transport remains to be studied.

The soluble sugars accumulation and carbohydrates partitioned to fruit are dramatically influenced by source and sink strength, the latter being the product of sink size and sink activity (Ho, 1988; Li et al., 2021). The dynamics of plant source-sink relationships are determined by key characteristics including photosynthetic carbon assimilation in source leaves, sucrose transport through the vasculature, type and strength of sink tissues, and metabolic status of sink tissues (Zhang et al., 2005; Jyotirmaya et al., 2021). Photosynthetic leaves serve as the primary source tissue organs in plants, with their morphological characteristics and photosynthetic intensity largely influencing the source strength (Mathan et al., 2016). Sink strength depends on the capacity to utilize photosynthates towards storage and maintenance while the utilization rate of photosynthetic products is directly affected by sink size, quantity, the functions of sugar metabolic enzymes (e.g., Cell wall invertase (CWINV) and sucrose synthase catalysis direction (SS-C)) and sugar re-synthases in fruits (SPS and SS-S) (Smith et al., 2018; Stein and Granot, 2019). Furthermore, the transport of photosynthetic products from source leaves to sink organs in the form of sucrose is a fundamental characteristic of plant source-sink balance, necessitating both symplastic and apoplastic phloem loading mechanism (Ruan, 2014). Previous studies have demonstrated that sucrose primarily moves from mesophyll cells of source leaves to parenchymal cells of phloem via plasmodesmata, establishing a symplastic pathway (Schulz, 2015; Miras et al., 2022). However, certain plants such as grape (Zhang et al., 2006), jujube (Nie et al., 2010), maize (Bezrutczyk et al., 2018), and potato (Abelenda et al., 2019) export sucrose from parenchymal cells to the apoplastic space through SWEET-mediated apoplastic pathways. Subsequently, it is actively absorbed into the sieve molecule-associated cell complex by sucrose/H^+^ symporter (SUTs) for transfer to sink organs (Chen et al., 2012; Ruan, 2014).

It has been suggested that K influenced cell swelling, with the cell pressure potential showing a positive correlation with fruit sugar content and hardness (Tong et al., 1999; Lester et al., 2006). Another perspective suggested that K affected photosynthesis in source leaves and promoted sugar metabolism and accumulation in storage organs (Lester et al., 2010), reducing K supply could decrease carbohydrate allocation to roots, resulting in an increased crown-to-root ratio (Ache et al., 2001). However, the underlying molecular mechanism by which K regulated the source-sink relationship to promote sugar accumulation in citrus remains elusive. We hypothesized that K influenced source and sink strength through its impact on carbohydrate distribution and transport processes within the plant system. Additionally, we proposed that K stimulated carbon flow between sources and sinks via the modulation of symplastic loading and/or apoplastic loading, thereby enhancing soluble sugars accumulation.

## Materials and Methods

### Experimental materials and design

A field experiment was conducted to study the effects of different amount of K fertilizer applications on soluble sugars accumulation and its mechanism by using 10-year-old Newhall navel orange (*Citrus sinensis*) grafed on trifoliate orange at the citrus research institute of Ganzhou, Jiangxi province in 2020. The soil properties are as follows: pH, 4.35; organic matter, 14.43 g·kg^-1^; alkaline hydrolysable N, 35.81 mg·kg^-1^; Olsen-extractable P, 73.63 mg·kg^-1^; exchangeable K, 183.76 mg·kg^-1^. The experiment was set up with five K fertilizer treatments including K0, K1, K2, K3 and K4, of which applied K (K_2_O) of 0, 0.25, 0.5, 0.75 and 0.9 kg·plant^-1^; N fertilizer (N): 0.8 kg·plant^-1^, P fertilizer (P_2_O_5_): 0.4 kg·plant^-1^, and provided with K_2_SO_4_, CO(NH_2_)_2_ and NH_4_H_2_PO_4_, respectively. The strategy of N, P and K fertilizer application followed our previous studies (Wu et al., 2021), namely bud fertilizer with 40 % N, 60 % P and 30 % K; Fruit preserving fertilizer with 30 % N, 40 % P and 50 % K; Fruit expanding fertilizer with 30 % N and 20 % K. The leaves and fruits were sampled at the stages of fruit enlarge (late Jul.), fruit trans-color (early Oct.) and fruit mature (late Nov.),and stored in -80°C for the analyses of carbohydrate, enzymes activity and genes expression.

To confirm the effects of K fertilizer application on fruit soluble sugar accumulation, a pot experiment were carried out in Huazhong Agricultural University, Wuhan, Hubei province by using 3-year-old Newhall navel orange grafed on trifoliate orange. The soil properties are as follows: pH, 6.62; organicmatter, 11.08 g·kg^-1^; alkaline hydrolysable N, 65.19 mg·kg^-1^; Olsen-extractable P, 18.82 mg·kg^-1^; exchangeable K, 68.78 mg·kg^-1^. The uniform orange seedlings were cultivated in a plastic pot with 35 kg soil. During the period of experiment, the pot were placed under a transparent rain proof shed and watered as needed with distilled water. The experiment was set up with four K levels of K0, K1, K2 and K3, and the application rates of K (K_2_O) were 0, 0.20, 0.40 and 0.60 g K_2_O kg^−1^ soil, and each treatment was replicated 5 times with 5 pots. The amount of N and P fertilizer were CO(NH_2_)_2_ (as N) 0.2 g·kg^−1^ soil and NH_4_H_2_PO_4_ (P_2_O_5_) 0.15 g·kg^−1^ soil, respectively. The strategy of N, P and K fertilizer was consistent with the field experiment. Fruits were collected at the mature stage for fruit quality analysis.

### Analysis of fruit quality

The fruit at the at each time period were selected to determined the fruit quality including total soluble solid (TSS), vitamin C (Vc). The concentration of TSS was determined by a digital refractometer, the vitamin C concentration was determined by 2,6-dichloroindophenol titration following the previous method (Wu et al., 2021).

### Analysis of soluble sugar and starch

The different stages of citrus leaves were ground with liquid nitrogen for the analysis of soluble sugar and starch. The homogenized samples of leaf were applied to extract soluble sugar and starch using 10 mL 80% (v/v) ethanol. Subsequently, the sample was centrifuged and the supernant were used to measure soluble sugar and starch according to the previous study (Ji et al., 2020).

### Photosynthetic data collection

Photosynthetic parameters (Transpiration rate (Tr); Net photosynthetic rate (Pn); Intercellular CO_2_ (Ci); Stomatal conductance (Gs)) were measured using the Li-6400 Portable Photosynthesis Measurement System (LI-COR, Lincoln, Nebraska, USA), from 8:00 to 12:00. The photosynthesis instrument was set as follows: the CO_2_ concentration of the air in the chamber was controlled by the carbon dioxide cylinders, while light used for the measurements was supplied by the LI-6400 LED red/blue light source, the gas flow rate, the mixing fan speed, the humidity and the air temperature of the leaf compartment were 10000 r/min, 500 µmol·s^-1^, 50 % and 25°C, respectively

### Analysis of sugar concentrations

The frozen samples of fruit were ground with liquid nitrogen for sugar concentrations determination including sucrose (Suc), fructose (Fru) and glucose (Glu) according to our previous studies (Wu et al., 2023). The homogenized samples were extracted by 80% methyl alcohol at 75°C for 15 min, then for ultrasonic extraction for 45 min and centrifuged at 4000×g for 10 min. After three times extractions, 0.50 mL supernatant were for drying by vacuum centrifugal concentration meter. After derivatization, 0.50 mL of supernatants were transferred into a 2 mL brown injection bottle and run-on Agilent 6890 N GC (Agilent, USA).

## 13C-isotope labeling and analysis

On the sunny side of the periphery of each tree crown, single fruit-bearing shoots with the same growth potential were selected for ring cutting and marked for feeding with ^13^CO_2_. Considering the effects of leaf-to-fruit ratio on leaf photosynthesis, all the fruit-bearing shoots were artificially adjusted to retain one marked mature leaves by picking leaves. The ^13^CO_2_ was supplied with NaH^13^CO_3_, one leaf was put into a plastic self sealing bag containing 0.11 g NaH^13^CO_3_, ^13^CO_2_ were released by injecting dilute H_2_SO_4_. After 6 h labeling, the fruit samples were harvested, and stored -80°C for the analysis of ^13^C-Suc, ^13^C-Fru and ^13^C-Glu. Following derivatization, The supernatants were used to analyzed ^13^C-Suc, ^13^C-Fru and ^13^C-Glu by using GC-IRMS.

### Micro-structure Observation and CFDA labeling

Plasmodesmata observations: The method of tissue preparation for ultrastructural observation was described by (Chen et al., 2017). Single-fruit branches were selected at the enlarge, trans-color and mature stages of citrus fruits, and the main leaf veins were cut transversely, and incubated in 4% (v/v) glutaraldehyde for plasmodesmata observations under a transmission electron microscope (Hitachi, Japan). Plasmodesmata were counted at all cell interfaces, including the interfaces between sieve element (SE)/ companion cell (CC), SE/ phloem parenchyma cells (PP), CC/PP and PP/PP in each selected field as described by Chen et al (2017).

Vascular bundle observations: Main leaf vein mentioned above were incubated in FAA fixative for vascular bundle observations. The samples were cut into segments and then were used to make paraffin section for the observations of vascular bundle development with a microscope (Nikon, DS-Ri2) according to the methods of Chen et al (2017). SE, sieve element; CC, companion cell; PP, phloem parenchyma cells; PD, plasmodesma CFDA labeling: To examine the Suc transport by symplastic pathway in the phloem of citrus, a feeding experiment was conducted using 5 (6)-carboxyfluorescein diacetate (CFDA) (Chen et al., 2017). Single-fruit branch was labeled with 200 µL of 1 mg·mL^-1^ CFDA aqueous solution. The citrus phloem was harvested after 48 h of in vivo transport of CFDA, which was selected for transverse and longitudinal cuts, sealed with 80% glycerol, and observed using a laser confocal microscope (Lecia SP8, Germany) with an excitation wavelength of 488 nm.

### Analysis of enzymes activities related to sugar metabolism

The 1 g fruit samples were homogenized in 9 mL PBS buffer (pH 7.2) on ice bath for 10 min, and then centrifuged at 3000 r·min^-1^ for 15 min at 4°C, and the supernatant were used for the determination of SPS, SS-S, SS-C and INV activities by using the ELISA Kits of these enzymes (Meibiai Biology, China) according to the manufacturer’s instructions.

### RNA Extraction and real-time PCR

Total RNA of leaf and fruit pulp was extracted using Hipure Plant total RNA kit (Magen Biotech, China) as described by the manufacturer’s protocol. After assessing the RNA quality and quantity, the 1.0 µg of high-purity RNA was taken to synthesize cDNA using TRUEscript RT kit (Aidlab, China). Real-Time PCR was performed using ABI 7500 (Bio-Rad, USA) according to the instructions of MonAmp^TM^ ChemoHS qPCR mix (Monad, China). Real-Time PCR conditions were as follows: 95°C, 10 min, followed by 40 cycles of 95°C for 10 s, 60°C for 30 s. The relative quantitative expressions were calculated by the 2^-ΔΔCt^ method, and the template primers were listed in Table S1.

### Statistical analysis

Statistical analyses of data were performed by SPSS 20.0 applying Duncan multiple comparison. The figures were drew using Origin 2018 software.

## Results

### K application improves fruit yield and quality in Newhall navel orange

The improvement of fruit enlargement and trans-color induced by K application were observed under pot culture, which is consistent with the results that K application increased the single fruit weigh, fruit number per plant and yield in Newhall navel orange under field culture (Figure. 1A-1D & Table S2). Compared with the control, K applications significantly increased the concentrations of soluble solid (TSS) of Newhall navel orange at mature stage, but no significant differences were observed in vitamin C (Vc) (Figure. 1E-1H) under pot and field culture. Compared with K0 treatment, the concentrations of K in leaf were significantly increased by K applications at the stages of enlargement and mature. Similarity, the concentrations of K in fruit pulp were dramatically enhanced by K applications at the stages of trans-color and mature (Figure S1). The result showed that appropriate levels of K application improved fruit quality in Newhall navel orange.

**Figure 1.**
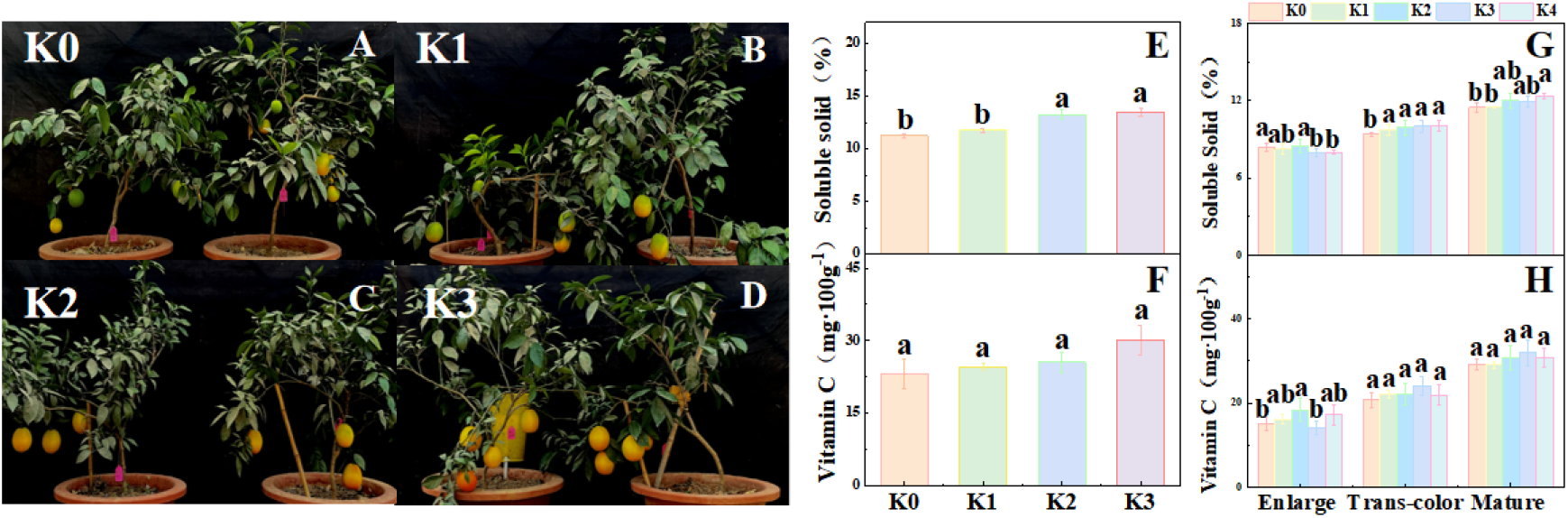
Effects of K fertilizer on fruit yield and quality in Newhall navel orange. (A-F) Pot culture: (E) soluble solids; (F) vitamin; K0, K1, K2 and K3 indicate the citrus plants were cultivated in different K fertilizer levels of 0, 0.20, 0.40 and 0.60 g K_2_O kg^−1^ soil under pot culture. (G-H) Field culture : (G) soluble solids; (H) vitamin; K0, K1, K2, K3 and K4 indicate that the citrus plants were cultivated in different K fertilizer levels of 0, 0.25, 0.5, 0.75, 0.9 kg K_2_O per plant under field culture. The data and error bars are the mean ± SE (n=4), and different lowercase letters represent significant differences among K treatments at same stage by Duncan-test (*P < 0.05*).

### K application improves the accumulations of soluble sugar in Newhall fruit

Suc, Fru, Glu are the main components of soluble sugar in citrus fruit (Wu et al., 2021), and we proved that the concentrations of Suc, Fru, Glu were significantly increased by K applications during the development of Newhall navel orange (Figure. 2). Compared with the control, the concentrations of Suc were dramatically increased at the stage of enlargement in K1, K2, K3 and K4 treatments with the increased rates of 20.38%, 22.31%, 31.08% and 38.52%, at the stage of trans-color in K2 and K3 treatments with increment of 8.40% and 9.19%, at the stage of mature in K3 and K4 treatments with increment of 6.11% and 5.02% in Newhall navel orange. The concentrations of Fru were significantly increased at the stage of enlargement in K1, K2 and K3 treatments with the increased rates of 8.46%, 9.08% and 8.71%, at the stage of mature in K1, K3 and K4 treatments with increment of 9.42%, 13.97% and 9.46%. The concentrations of Glu were markedly enhanced by 7.95%, 8.34%, and 10.94% at the stage of enlargement in K1, K2 and K4 treatments, by 11.33%, 10.39%, 22.73%, 14.82% at the stage of mature in K1, K2, K3 and K4 treatments in Newhall navel orange (Figure 2). To further elaborate the relationship of fruit sugar concentrations and K concentrations of pulp, we performed the regression model between sugar and K concentrations. We found that Suc concentrations was significantly correlated with K concentrations of pulp (Figure S3). Moreover, 1.50% of K concentration in pulp was the threshold value of K concentration for the decrease of Suc in fruit that caused by excessive K.

**Figure 2.**
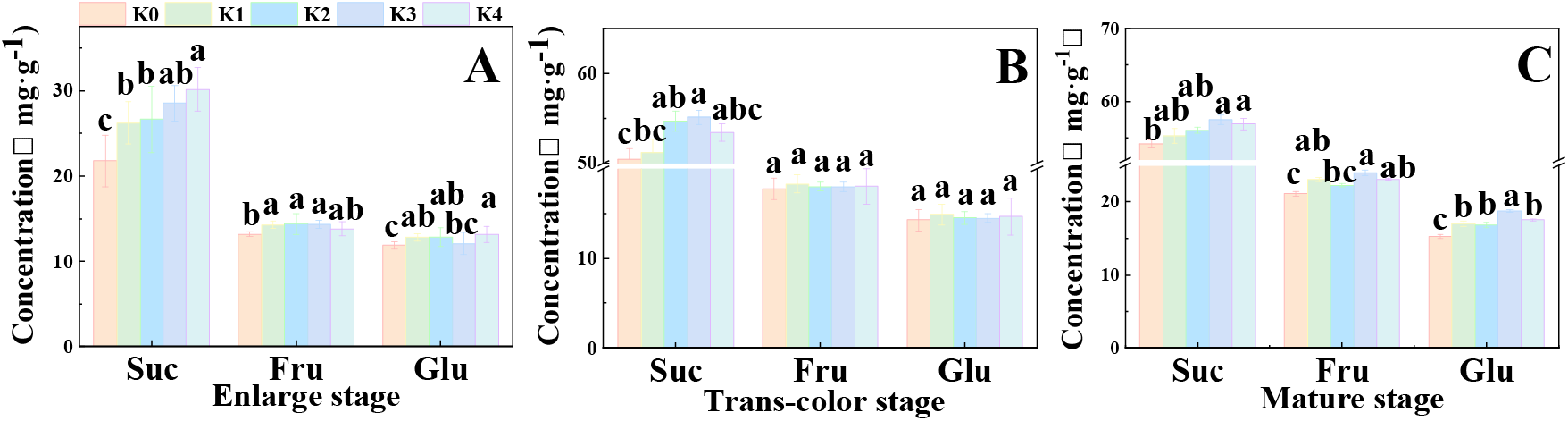
Effects of K fertilizer on soluble sugar concentrations in Newhall navel orange fruit. (A) enlarge stage; (B) trans-color stage; (C) mature stage. K0, K1, K2, K3 and K4 indicate that the citrus plants were cultivated in different K fertilizer levels of 0, 0.25, 0.5, 0.75, 0.9 kg K_2_O per plant under field culture. Suc, Fru and Glu represent sucrose, fructose and glucose, respectively. The data and error bars are the mean ± SE (n=4), and different lowercase letters represent significant differences among K treatments at same stage by Duncan-test (*P < 0.05*).

### K application improves Suc metabolism and sink strength in Newhall fruit

The Suc metabolism of fruit itself affects Suc synthesis and sink strength in fruit (Julius et al., 2017), we first investigated the effects of K application on Suc metabolism. Compared with K0 treatment, K2 and K4 treatments increased Suc-synthesizing enzymes SPS and SS-S activities, and significant increases were observed in SPS activity of K2 and K4-cultivated fruit at the stage of enlargement, in SS-S activity of K4-cultivated fruit at the stage of enlargement, and in SPS and SS-S activities of K2-cultivated fruit at the stage of trans-color (Figure 3A & 3B). Compared with K0 treatment, K4 treatment significantly enhanced Suc-degrading enzyme SS-C activity at the stage of enlargement (Figure 3C). However, the activities of SS-C and INV were dramatically inhibited by K4 at the stage of trans-color (Figure 3C & 3D).

**Figure 3.**
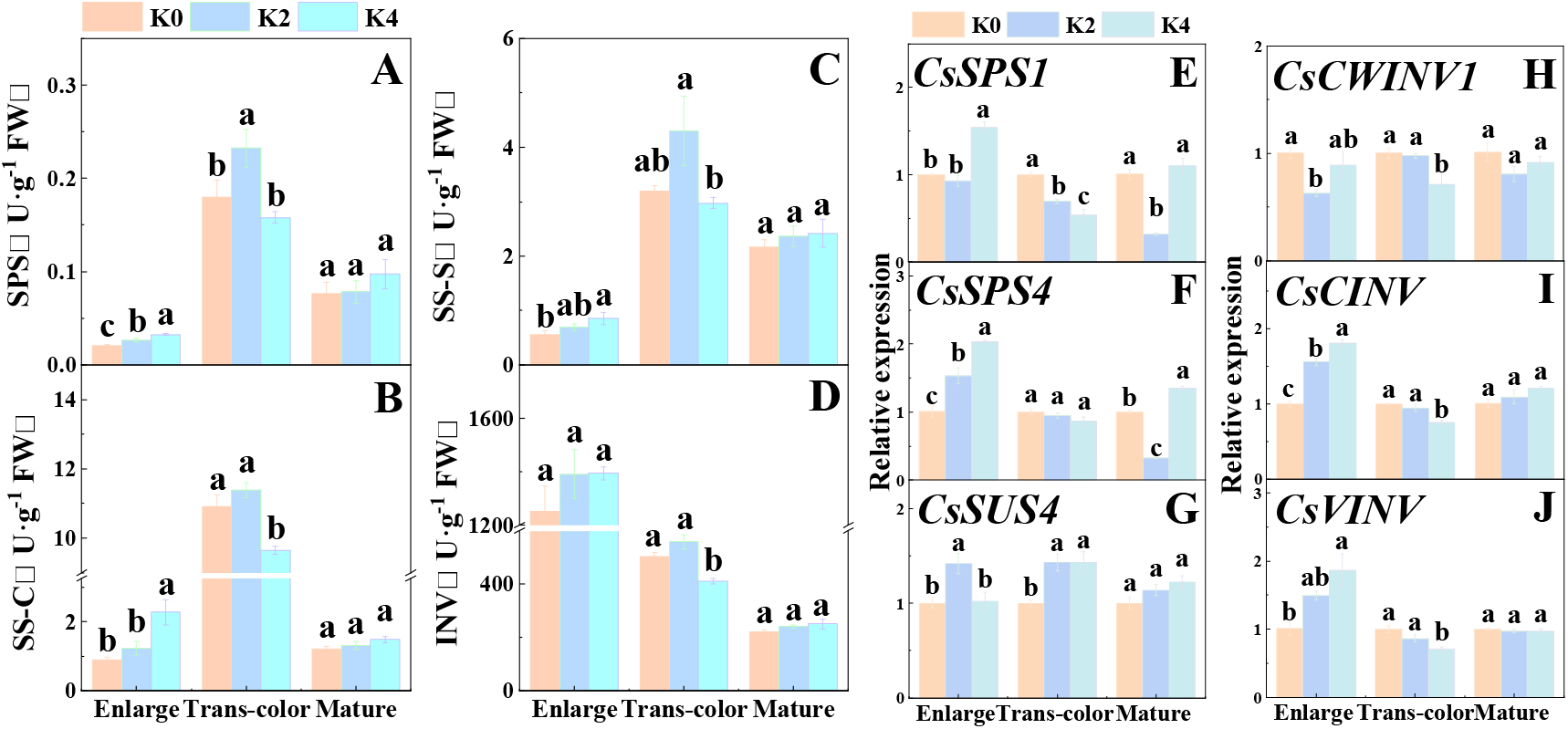
Effects of K fertilizer on the activity and gene expression of Suc-metabolizing enzymes in Newhall navel orange fruit. (A) sucrose phosphate synthase (SPS); (B) Sucrose synthetase-cleavage (SS-C); (C) Sucrose synthetase-synthesis (SS-S); (D) Invertase (INV). K0, K2 and K4 indicate that the citrus plants were cultivated in different K fertilizer levels of 0, 0.5, 0.9 kg K_2_O per plant under field culture. The data and error bars are the mean ± SE (n=4), and different lowercase letters represent significant differences among K treatments at same stage by Duncan-test (*P < 0.05*).

Compared with K0 treatment, the expressions of *CsSPS1* and *CsSPS4* were significantly increased by K2 and/or K4 treatment at enlargement stage, but the expressions of *CsSPS1* and *CsSPS4* were markedly decreased by K2 treatment at mature stage (Figure 3E & 3F). Similarly, the expressions of *CsSUS4* were significantly increased by K2 and/or K4 treatment at enlargement and trans-color stages (Figure 3G). Compared with K0 treatment, the expressions of *CsCINV* and *CsVINV* were markedly increased by K2 and/or K4 treatment at enlargement stage, but the expression the expressions of *CsCWINV1, CsCINV* and *CsVINV* were observably decreased by K4 treatment at trans-color stage.

### K application improves the transport of sugar in in Newhall fruit

Compared with K0 treatment, the expressions of *CsSUT1*, *CsSUT2*, *CsSUT4*, *CsSTP11*, *CsSWEET16* and *CsTMT2* were significantly increased by K2 and K4 applications, and the expressions of *CsSWEET15* and *CsVGT2* were dramatically induced by K2 at enlargement stage (Figure 4). Correspondingly, the significant increases of *CsSUT1*, *CsSUT2* and *CsTMT2* expressions were observed under K2 and K4 treatment at trans-color stage, and the increased expressions of *CsSUT4* and *CsSWEET15* occurred in K2 treatment at trans-color stage.

**Figure 4.**
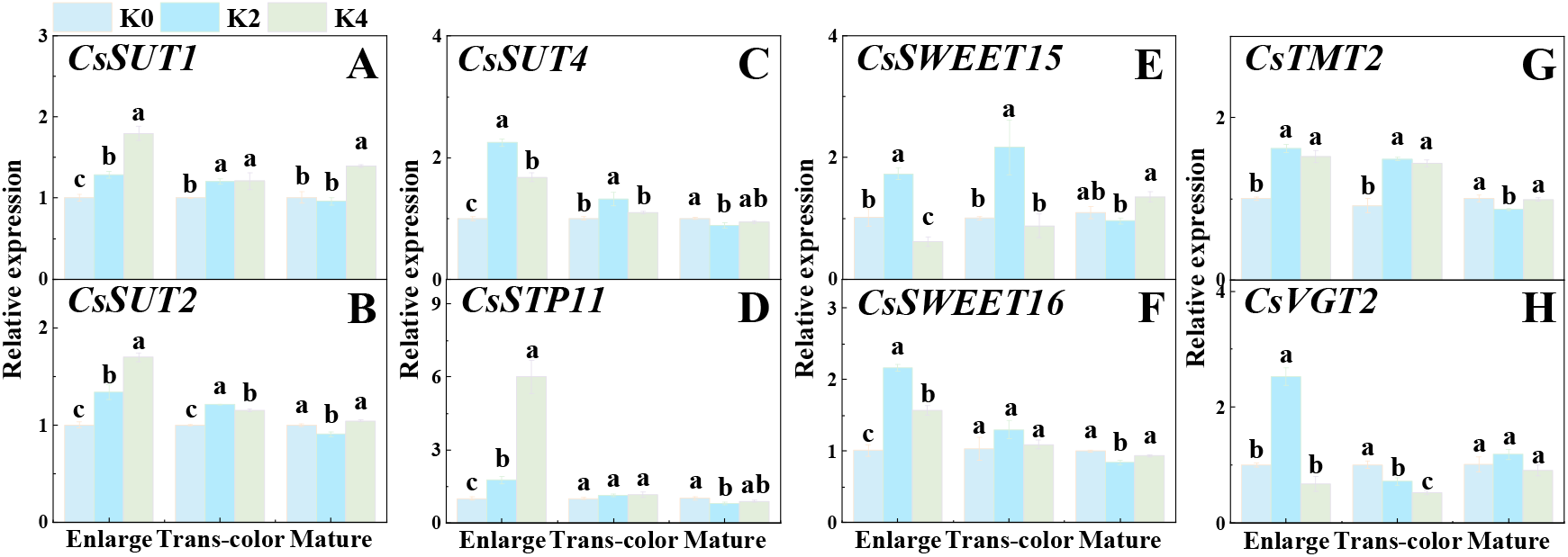
Effects of K fertilizer on the expressions of sugar transporters in Newhall navel orange fruit. K0, K2 and K4 indicate that the citrus plants were cultivated in different K fertilizer levels of 0, 0.5, 0.9 kg K_2_O per plant under field culture. The data and error bars are the mean ± SE (n=4), and different lowercase letters represent significant differences among K treatments at same stage by Duncan-test (*P < 0.05*).

### K application improves sugar metabolism and source strength in Newhall leaf

Source strength determines the amount of carbohydrate export (Lemoine et al., 2013). To further understand the cause of the increase of soluble sugar, we measured the effects of K application on leaf sugar metabolism. Compared with K0 treatment, the transpiration rate (*Tr*) and stomatal conductance (*Gs*) of Newhall leaves were sharply enhanced by K1 and K2 treatments. Similarity, the net photosynthetic rate (*Pn*) of Newhall leaves were significantly increased by K applications with increment of 43.08%∼84.57% (Table S3), indicating that K fertilizer can improve the fixation of carbon. Compared with K0 treatment, the expressions of *CsSPS2*, *CsSPS3* and *CsSPS4* were significantly up-regulated under K2 and K4 treatments at enlargement and trans-color stages, but the expressions of *CsCWINV1*, *CsCINV* and *CsVINV* was significantly down-regulated at enlargement and mature stages (Figure 5). The result indicated that K application improved the synthesis of Suc in Newhall leaves at the stages of enlargement and mature. However, no significant differences were observed on soluble sugar and starch concentrations among K treatments during the period of fruit development, indicating that K applications may promote Suc transport from leaf (source) to fruit (sink).

**Figure 5.**
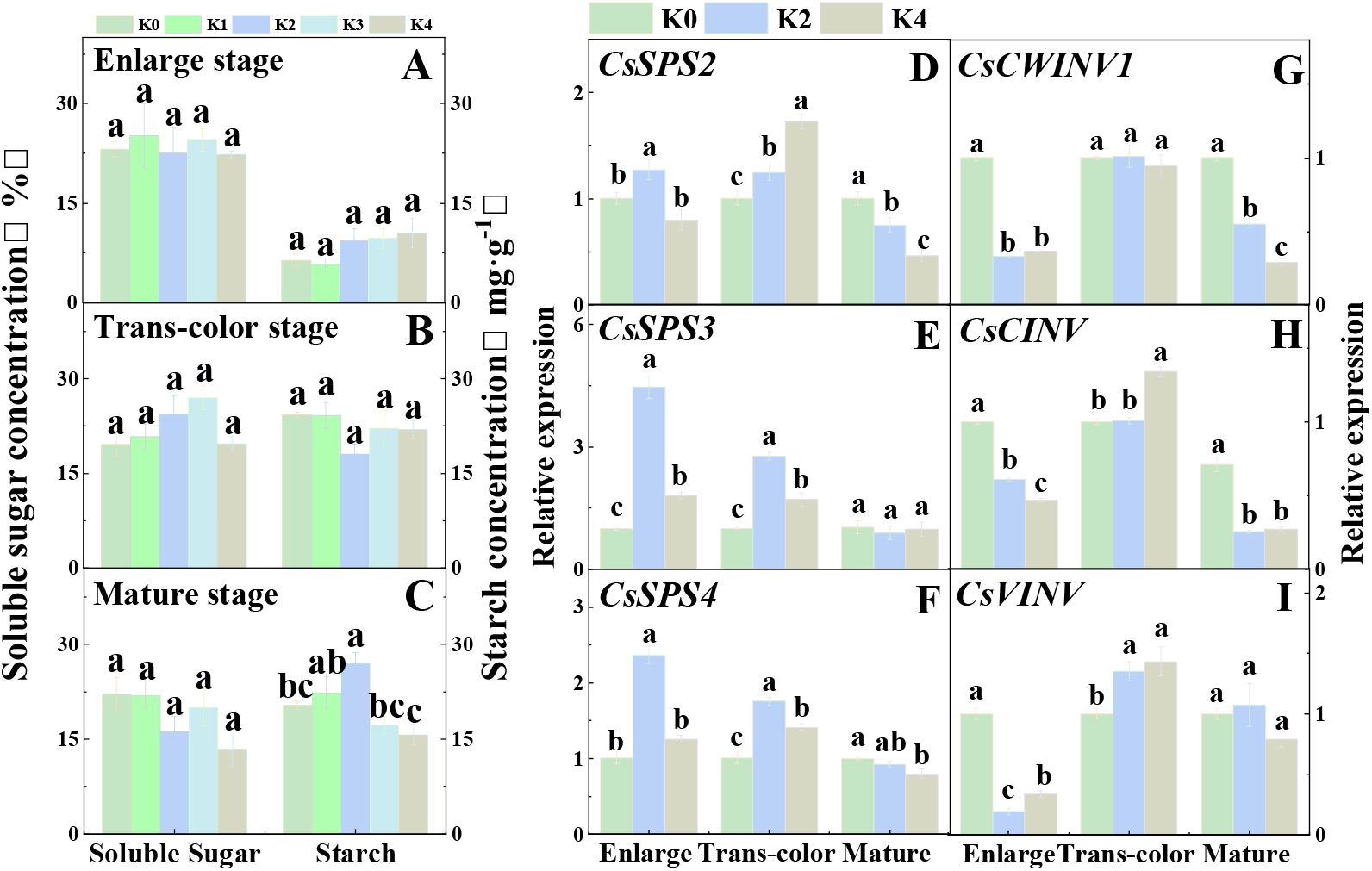
Effects of K fertilizer on the concentrations of soluble sugar and starch and the expressions of Suc-metabolizing enzyme gene in Newhall leaf. K0, K1, K2, K3 and K4 indicate that the citrus plants were cultivated in different K fertilizer levels of 0, 0.25, 0.5, 0.75, 0.9 kg K_2_O per plant under field culture. The data and error bars are the mean ± SE (n=4), and different lowercase letters represent significant differences among K treatments at same stage by Duncan-test (*P < 0.05*).

### K application improves carbohydrate transport from source to sink in Newhall

Carbohydrate transport plays important roles in the accumulations of soluble sugar in fruit (Sadka et al., 2019) and half of the sugars in citrus fruit come from carbohydrate transport (Koch, 1984). To test the roles of K in regulating carbohydrate transport in citrus, we performed ^13^C-isotope experiments (Figure 6A). The results showed that K applications significantly increased the concentrations of ^13^C-Suc in Newhall fruit at enlargement stage and the concentrations of ^13^C-Suc and ^13^C-Glu in Newhall fruit at trans-color stage, suggesting that K promoted sugar transport from source to sink in Newhall (Figure 6B).

**Figure 6.**
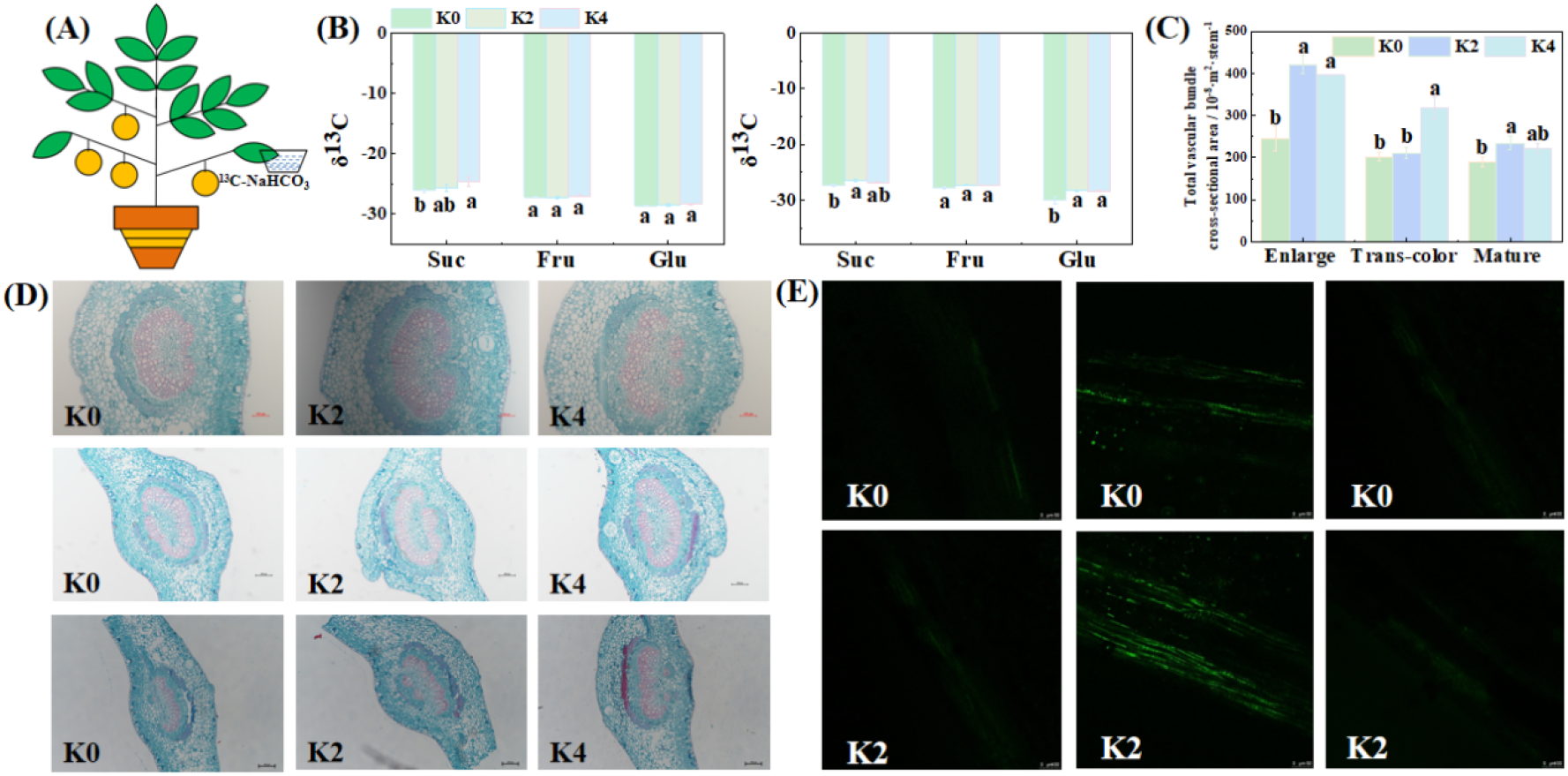
Effects of K fertilizer on phloem transport of carbon flow in Newhall leaf-to-fruit. K0, K2 and K4 indicate that the citrus plants were cultivated in different K fertilizer levels of 0, 0.5, 0.9 kg K_2_O per plant under field culture. (A) ^13^C-NaHCO_3_ labeling experimental model. (B) ^13^C-sugar components in fruit. (C) the total cross-sectional area of vascular bundle of leaf vein. (D) Cross-sectional sections of vascular bundles of leaf veins. (E) CF fluorescence signal in phloem. The data and error bars are the mean ± SE (n=4), and different lowercase letters represent significant differences among K treatments at same stage by Duncan-test (*P < 0.05*).

Carbohydrate transport in phloem loading mainly relies on two kinds of pathway, namely symplastic and apoplastic pathway (Ruan, 2014), we firstly evaluated the roles of K on carbohydrate transport that by symplastic pathway. Compared with K0 treatment, K applications increased the total cross-sectional area of vascular bundle at different stages, significant increases were observed in K2 and K4 treatments at the stage of enlargement, in K4 treatment at the stage of trans-color and in K2 treatment at the stage of mature (Figure 6C & 6D). Further, we used symplastic tracer carboxy fluorescein diacetate (CFDA) to confirm the symplastic pathway of carbon flow in phloem. CFDA was fed into branches far from fruit, where it was cleaved into CF that only diffuses through plasmodesmata (Nie et al., 2010). The results showed that CF signals were strengthened by K applications, particularly the strongest signal were observed at the stage of trans-color (Figure 6E). Interestingly, the plasmodesmal densities (PD) of SE/CC, SE/PP, PP/PP were increased by K applications, and higher PD of SE/CC and PP/PP were observed at the stage of trans-color, which is consistent with the results of CF signals in phloem in Newhall orange (Figures 7, S4, S5 & Table S4). These results indicated that K applications improved the carbon allocation from source to sink in Newhall fruit, and the transport of carbon flow at trans-color stage was a critical stage for phloem transport by symplastic pathway.

**Figure 7.**
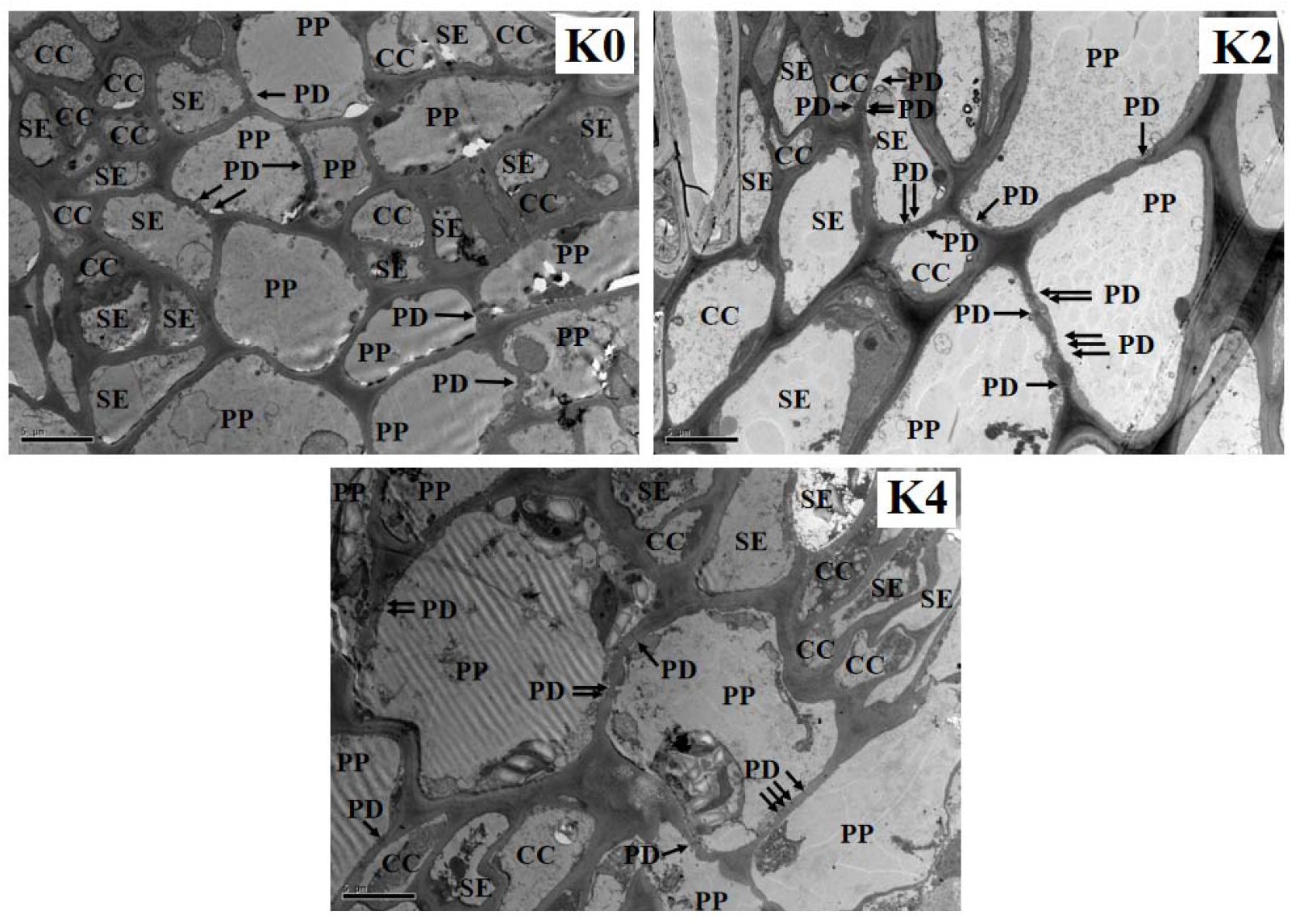
Ultrastructure of the sieve element-companion cell complex and its surrounding phloem parenchyma cells of Newhall orange leaf vein in the trans-color stage. SE, sieve element; CC, companion cell; PP, phloem parenchyma cells; PD, Plasmodesma. K0, K2 and K4 indicate that the citrus plants were cultivated in different K fertilizer levels of 0, 0.5, 0.9 kg K_2_O per plant under field culture.

Apoplastic pathway was the another pathway that carbon flow transport in phloem, the transporters of SUTs and SWEETs are responsible for the phloem loading (Scofield et al., 2002; Yang et al., 2018; Wang et al., 2021), we further evaluated the roles of K on carbohydrate transport that by apoplastic pathway. Compared with K0 treatment, the expressions of *CsSUT1*, *CsSUT2*, were significantly increased under K2 and K4 treatment at enlargement stage, but the expressions of *CsSWEET16* were decreased (Figure 8). K applications increased the expressions of *CsSUT1*, *CsSWEET16* at trans-color stage, significant increases occurred in K2 treatment of *CSSUT1* and in K2 and K4 treatments of *CsSWEET16*. These results indicated that the increased expressions of *CsSUT1*, *CsSWEET16* caused by K may also involved in the regulation of carbon flow transport from leaf to fruit at the enlargement and trans-color stages.

**Figure 8.**
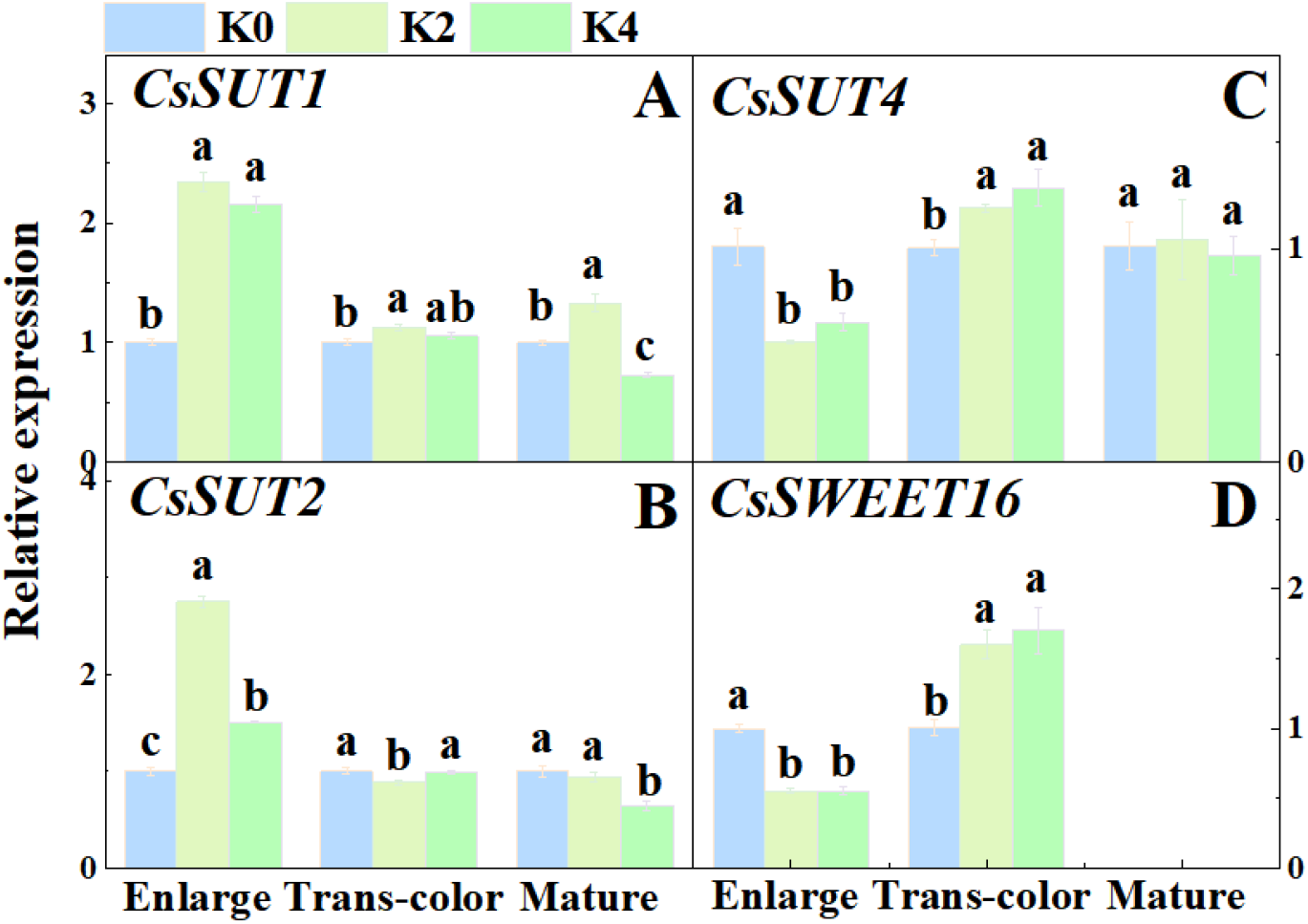
Effects of K fertilizer on the expressions of sugar transporter in Newhall leaf. K0, K2 and K4 indicate that the citrus plants were cultivated in different K fertilizer levels of 0, 0.5, 0.9 kg K_2_O per plant under field culture. The data and error bars are the mean ± SE (n=4), and different lowercase letters represent significant differences among K treatments at same stage by Duncan-test (*P < 0.05*).

## Discussion

### Appropriate K concentration improved fruit quality and the accumulations of sugar in Newhall

K is an essential macro-nutrient in higher plants and also play crucial roles in fruit quality of perennial fruit trees (Wu et al., 2021). As previously reported, K application increased soluble sugar accumulations in apple (Zhang et al., 2018), pear (Shen et al., 2017), tomato (Schwarz et al., 2013) and grape (Niu et al., 2008). Although the primary phytochemical compounds with variable changes among fruits of different species, K fertilizer also increased the concentrations of soluble solids, Suc, Fru and Glu in Newhall navel orange (Figures 1 and 2), suggesting that K fertilizer may induced a series of changes that related to sugar accumulations. In citrus species, fruit sugar mainly consists of three major soluble sugars, namely Suc, Glu and Fru (Ladaniya, 2008). Moreover, the concentrations of soluble sugar in fruit are an important factor to determine its fruit quality (Zhou et al., 2018; Wu et al., 2021). Therefore, the increased concentrations of soluble sugar caused by K application will contributed to improving fruit quality in navel orange. However, excessive K application decreased the sugar concentrations and fruit quality (Wu et al. 2023). Generally, appropriate K supply is necessary to ensure plant growth and yield. For field crops, the concentrations of K in plant tissues are 2%-10% of dry matter (Leigh and Jones, 1984). However, the appropriate concentrations of K in fruit are still to be quantified. To meet with this gap, we found that 1.5% of K concentration was the optimal concentration by using regression model (Figure S3), moreover, higher this concentration will be detrimental to sugar accumulation and fruit quality formation. Taken together, appropriate K concentration improved fruit quality and the accumulations of sugar in Newhall fruit.

### The increased strength of sink and source caused by K contributed to increasing sugar accumulations in Newhall fruit

Sink strength and source strength regulate carbon allocation and play key roles in regulating crops productivity and horticultural plants fruit quality (Yu et al., 2015; Wu et al., 2021). Sink strength is the inherent ability of organs to absorb assimilates, which can be quantitatively expressed as the growth rate of organs (Marcelis, 1996). Therefore, the sink strength of single fruit (sink size and sink activity) and fruit number conjointly determine the total fruit sink strength, thereby affecting the proportion of carbon allocation to the fruits (MarcelisA, 1994; Ji et al., 2020). In the present study, we found that K application increased fruit size in K2 and K3 treatments and fruit number in K4 treatment (Table S2). Here, we did not compare the dynamic growth rate of sink organ (fruit) between K treatments, but the average growth rate of sink organ in K treatments was higher than it in no K application during the whole stage (Table S2), which was consistent with the result that the increased growth rate of tomato fruit caused by far-red radiation contributed to the improvement of sink strength of individual fruit (Ji et al., 2020). Early flowering is a response to shading, which may lead to an increase of fruit number and thus increase the sink strength (Yuan et al., 2017). However, appropriate K application induced a minor increase in fruit numbers (Table S2), indicating that the effects of K on fruit number did not contribute obviously to the increase of sink strength, which was lined with the results that far-red radiation increased sink strength was not depend on the increases of fruit number (Ji et al., 2020). Therefore, the results suggest that K application increases the sink strength of fruit by increasing growth rate, thus contributing to attraction of photoassimilates into fruit and hence improves sugar accumulations and yields in fruit (Figure 2; Table S2).

However, how K application improved sink strength of fruit? Sink strength regulated by Suc metabolism, and the activities of Suc-metabolizing enzyme, such as SPS, SS-S, SS-C and INV, play important roles in regulations of sink strength (Li et al., 2021). K application increased the activities of SPS, SS-S, SS-C and INV, the expressions of *CsSPS1*, *CsSPS4* and *CsSUS4* at the stages of enlargement and tans-color (Figure 3), which were supported by previous studies (Zahoor et al., 2017; Wu et al., 2021; Wu et al., 2023). Soluble sugars are degraded, re-synthesized, transported and eventually accumulated in the fruit organs, where cytoplasmic Suc is hydrolyzed by SS-C or INV to hexose for glycolysis or re-synthesized into Suc by SPS and SS-S (Nguyen-Quoc and Foyer, 2001). The increased activities of SS-C and INV at the stages of enlargement and tans-color, and the increased expressions of *CsCINV* and *CsVINV* at the stages of enlargement in K treatments will contributed to the degradation of the imported Suc from leaf and thus produced a steep of Suc concentration differences. In accordance with our findings in tomato (Wu et al. 2023), one part of the degraded products were re-synthesized into Suc due to the increased activities of SPS and SS-S caused by K application in citrus, another part of the degraded products were used for energy metabolism, carbon skeleton, and sugar signal (Ruan, 2014). Sink strength also regulated by sugar transport, and thus sugar transporters play important roles (Yu et al., 2015; Ji et al., 2020). We found that K increased the expressions of *CsSWEET15* in newhall fruit, a gene is responsible for sugar unloading in phloem, and thus enhanced Suc efflux from phloem, which was supported by a study in tomato (Ko et al., 2021). Subsequently, the degraded products of Suc, namely Fru and Glu, were transported into vacuole for storage by *CsSTPs*, *CsTMTs* and *CsVGTs* (Zheng et al., 2014). Therefore, the increased expressions of *CsSTP11*, *CsTMT2* caused by K at the stages of enlargement and tans-color will drive the sugars into vacuole and hence increased the sink capacity of fruit to attract more photoassimilates into fruit at the early and mid stages of fruit development. Taken together, K application improved sink strength by regulating sugar metabolism and transport and thus will contribute to increasing sugar accumulations in Newhall fruit.

Source strength determines the amount of carbohydrate export (Lemoine et al., 2013). Unlike sink strength of fruit, source strength of leaf is not easily measured through experiments, while the fixation capacity of carbon and the synthesis capacity of carbohydrates can reflect the source activity (Ho, 1996; Lemoine et al., 2013; Yu et al., 2015). The present study showed that K application increased photosynthesis (*P_n_*) (Table S3), indicating that K improved carbon fixation and thus stimulated source activity. Source strength can also be reflected through Suc synthesis activity in leaf (Yu et al., 2015). K application increased the expressions of Suc-synthesized enzyme gene *CsSPS2*, *CsSPS3*, *CsSPS4* of leaf at the stages of enlargement and tans-color, but decreased the expressions of Suc-degraded enzyme gene *CsCWINV1*, *CsCINV* at the stage of enlargement, indicating that source organ can supply more Suc to export into sink organ in K treatments due to the increased Suc synthesis and the decreased Suc degradation, and make a steep of Suc concentration differences to drive carbon flow between source and sink in Newhall orange.

### 4.3 K stimulates carbon flow between source and sink and hence increases fruit sugar in Newhall

Previous study showed that more than half of fruit sugars are from source leaf through the phloem transport in citrus fruit, suggesting that carbohydrates transport play crucial roles in fruit sugar accumulation (Koch, 1984; Katz et al., 2011). We have previously confirmed that K can enhance the strength of the source and sink (Figure 3), meaning source leaf can provide more carbohydrates, while the sink can provide a larger storage space, which creates a prerequisite for the carbon flow between source and sink. However, whether and how K regulates carbohydrates transport from source to sink still to be studied. Although K increased carbon fixation and source activity, the concentrations of leaf soluble sugar were not increased by K treatments (Figure 5). Starch was a main synthetic product when soluble sugar concentrations was increased in source leaf (Nebauer et al., 2011), but the the concentrations of leaf starch were not enhanced under K-treated condition (Figure 5), even the previous study showed that K improved the activities of starch-synthesized enzyme (Gao et al., 2021). Therefore, it was implied that the increased carbohydrates were transported into fruit, and this confirmed by ^13^C-isotope experiments that the concentrations of ^13^C-Suc and ^13^C-Glu were increased by K application (Figure 6B), which indicated that K promoted sugar transport from source to sink in Newhall.

However, how K regulates carbohydrates transport from source to sink? The carbohydrates are firstly loaded into sieve element-companion cell complex (SE-CC) in leaf phloem, transported over a long distance to the fruit, and then unloaded on the segment epidermis of fruit by symplastic or apoplastic pathway (Zhang and Turgeon, 2018; Paniagua et al., 2021). The symplastic pathway of carbohydrates entering the companion cell mainly passed through plasmodesmata, and the resistance of this pathway is relatively low (Hu et al., 2011). We found that the phloem loading of carbohydrates can use the symplastic pathway in Newhall (Figure 6E), which was reflected by a symplastic tracer of CFDA (Nie et al., 2010). Moreover, the CF signal intensity at tans-color stage was higher than that at enlargement and mature stages, interestingly, K improved CF signal intensity at tans-color stage, indicating that symplastic loading of carbohydrates at tans-color stage play more important roles than that at enlargement and mature stages. Symplastic loading is a plasmodesmata-mediated process, the PD thus is very important to appraise its transport capacity (Hu et al., 2011; Ma et al., 2019; Sun et al., 2019). In accordance with the result of symplastic tracer (Figure 6E), the PDs of SE/CC, CC/PP and PP/PP at tans-color stage were higher than those at enlargement and mature stages, importantly, K increased the PD of SE/CC at the whole stages in K2 treatment (Figures 7, S3-S4; Table S4), which further verified that phloem loading of carbohydrates is mainly carried out through symplastic pathway at tans-color stage, and this process were strengthened by K application.

Besides, apoplastic loading is a transporters-mediated process, the SUTs and SWEETs transporters are responsible for evaluation of its transport capacity (Zhang and Turgeon, 2018; Ma et al., 2019). The increased expressions of *CmSUT1* and *CsSUT2* improved Suc concentrations in phloem sap and ^14^C containing-Suc concentrations in melon and cucumber, respectively (Gil et al., 2011; Ma et al., 2019). Drastic decrease of ^11^C-photoassimilate export from leaf of maize *sut1* mutant were observed by (Babst et al., 2022). The increased expressions of phloem loading gene *CsSUT1* and *CsSUT2* at enlargement and tans-color stages caused by K were observed (Figure 8), indicating that the apoplastic loading maybe improved by K, which were consistent with the findings in tomato (Wu et al. 2023). Interestingly, the expressions of *CsSUT1* and *CsSUT2* at enlargement stage were higher than those at tans-color and mature stages (Figure 8), suggesting that apoplastic loading of carbohydrates at enlargement stage play more important roles than that at tans-color and mature stages. Combined with the symplastic data, we concluded that the enlargement and tans-color stages were the key stages for carbon flow from source to sink, and apoplastic loading was the main route at enlargement stage, but symplastic loading was the main route at tans-color stage.

After phloem loading, the carbohydrates will transport over a long distance to the fruit, and transport capacity is controlled by flow resistance, which mainly depends on the anatomical structure of phloem and the turgor pressure-mediated driving force (Epron et al., 2016; Liesche, 2016). Epron et al (2016) showed that K increased carbon transport velocity in trunk phloem due to the increasing cross-sectional area of sieve elements in eucalyptus tree. In line with our findings, K increased cross-sectional area of vascular bundle of main vein. Moreover, the concentrations of phloem K were increased in Newhall branches by K application (Figure S2), which were consistent with the results in phloem sap of *Eucalyptus grandis* (Battie-Laclau et al., 2014). Importantly, the increased concentrations of K in phloem was conducive to maintaining phloem pressure and thus produced a driving force for carbon flow (Babst et al., 2022). Therefore, the cross-sectional area of vascular bundle and K concentrations contributed to driving carbon flow in K-treated plants, but the carbon transport velocity inside phloem was further needed to measure in future work. Subsequently, the carbohydrates will be unloading on segment epidermis of fruit mainly by apoplastic pathway due to the higher expressions of *CsSWEET15*, *CsSWEET16* (Figure 4) and less plasmodesmata (unpublished data).

## Conclusions

Collectively, differences in enzyme activities of sugars metabolism, vascular features and sugar transporter functions led to a differential source-sink relationship upon different K application rates in citrus. K application induced source-sink dynamics favoring high sugar concentrations through the accumulation of carbohydrates, mediated by higher photosynthesis and expressions of *CsSPSs* in source leaves, higher SPS and SS activity and expressions together with higher expression of genes related vacuole sugar transport in fruit. Additionally, K application stimulated carbon flow through symplastic and apoplastic pathways, which were supported by the structural characteristics of the petiole and phloem cells and the expression of *CsSUTs* and *CsSWEETs*, ultimately promoting sugar accumulation. Taken together, this study elucidated that K application regulated source-sink relationship by enhancing source strength, promoting sink activity, and increasing carbon flow to promote sugar accumulation. Future research should focus on elucidating the precise contribution rate of K application in enhancing fruit sugar concentration by augmenting source strength, sink capacity, carbon flow and increasing metabolic flux of core metabolites in fruits.

## ACKNOWLEDGEMENTS

This research was supported by the Fundamental Research Funds for the Central Universities (2662022ZHQD002), the National Natural Science Foundation of China (No. 32001986), the National Key Research and Development Program of China (2019YFD1000103), the China Post-doctoral Science Foundation (No. 2021T140245) and the Modern Citrus Industry Technology System of China (CARS-26).

## AUTHOR CONTRIBUTION

Kongjie Wu: Investigation, Methodology, Data curation, Visualization, Writing-original draft. Chengxiao Hu: Supervision, Resources. Peiyu Liao: Investigation. Yinlong Hu: Investigation. Xuecheng Sun: Writing-review & editing. Qiling Tan: Data curation. Zhiyong Pan: Writing-review & editing. Shoujun Xu: Writing-review & editing. Zhihao Dong: Investigation. Songwei Wu: Funding acquisition, Validation, Resources, Supervision, Writing-review & editing.

## FUNDING STATEMEENT

This research was supported by the Fundamental Research Funds for the Central Universities (No. 2662022ZHQD002), the National Natural Science Foundation of China (No. 32001986), the National Key Research and Development Program of China (No. 2019YFD1000103), the China Post-doctoral Science Foundation (No. 2021T140245) and the Modern Citrus Industry Technology System of China (No. CARS-26).

## DATA AVAILABILITY STATEMENT

Data will be made available on request.

